# Cytosine Variant Calling with High-throughput Nanopore Sequencing

**DOI:** 10.1101/047134

**Authors:** Arthur C. Rand, Miten Jain, Jordan Eizenga, Audrey Musselman-Brown, Hugh E. Olsen, Mark Akeson, Benedict Paten

## Abstract

Chemical modifications to DNA regulate cellular state and function. The Oxford Nanopore MinION is a portable single-molecule DNA sequencer that can sequence long fragments of genomic DNA. Here we show that the MinION can be used to detect and map two chemical modifications cytosine, 5-methylcytosine and 5-hydroxymethylcytosine. We present a probabilistic method that enables expansion of the nucleotide alphabet to include bases containing chemical modifications. Our results on synthetic DNA show that individual cytosine base modifications can be classified with accuracy up to 95% in a three-way comparison and 98% in a two-way comparison.

**Statement of Significance:** Nanopore-based sequencing technology can produce long reads from unamplified genomic DNA, potentially allowing the characterization of chemical modifications and non-canonical DNA nucleotides as they occur in the cell. As the throughput of nanopore sequencers improves, simultaneous detection of multiple epigenetic modifications to cytosines will become an important capability of these devices. Here we present a statistical model that allows the Oxford Nanopore Technologies MinION to be used for detecting chemical modifications to cytosine using standard DNA preparation and sequencing techniques. Our method is based on modeling the ionic current due to DNA k-mers with a variable-order hidden Markov model where the emissions are distributed according to a hierarchical Dirichlet process mixture of normal distributions. This method provides a principled way to expand the nucleotide alphabet to allow for variant calling of modified bases.

## Introduction

Eukaryotic DNA chemical modifications of cytosine (C) include 5-methylcytosine (5-mC), hydroxymethylcytosine (5-hmC), 5-formylcytosine, and 5-carboxylcytosine. DNA methylation is involved in multiple facets of biology, such as gene regulation, cell differentiation and development, and disease. In addition, 5-mC and 6-methyladenine (6-mA) are involved in bacterial gene regulation^1–3^.

Next generation sequencing technologies use chemical treatment to detect cytosine methylation. The treatment causes base substitutions that can be read without an expanded nucleotide alphabet. These techniques are limited by sequence read length of 100-500 base pairs and can only detect one cytosine variant at a time. Single-molecule real time (SMRT) sequencing generates long reads (1-5 kb), and researchers have shown that it can detect multiple modifications to DNA simultaneously using enzyme kinetics^4,5^. The Oxford Nanopore Technologies’ (ONT) MinION is a portable, low-cost single molecule DNA sequencer that can sequence long fragments of (50 kb) DNA at up to 92% accuracy absent amplification^6^.

Computational analysis of nanopore data has historically been a niche area of bioinformatic research^7,8^, but the field has broadened since the beginning of MinION Access Program in 2014. Recently published algorithms have focused on alignment and *de novo* genome assembly using hidden Markov models^9–12^. We build on this literature by taking a similar approach to detecting base modifications. Our group and others have previously shown that ionic current measurements from low-throughput nanopore sensors can discriminate all five C5-cytosine variants^13,14^. In this paper, we demonstrate that the MinION nanopore sequencer can discriminate among C, 5-mC, and 5-hmC at high-throughput without special DNA preparation.

Our method is based on a generative model of the MinION’s ionic current signal. In particular, we assume that the signal is emitted by a variable-order pair hidden Markov model (HMM) that tracks a reference sequence but allows a reference nucleotide to match any of several modified bases (Figure 1A-B, Figure 4). We augment the HMM by modeling the ionic current distributions with a hierarchical Dirichlet process mixture model (HDP), a Bayesian nonparametric method that shares statistical strength to robustly estimate a set of potentially complex distributions^15^. We show that the HDP meaningfully enhances the HMM’s ability to detect cytosine variants by comparing it to a simpler HMM with emissions modeled by parametric normal distributions.

**Figure 1.**
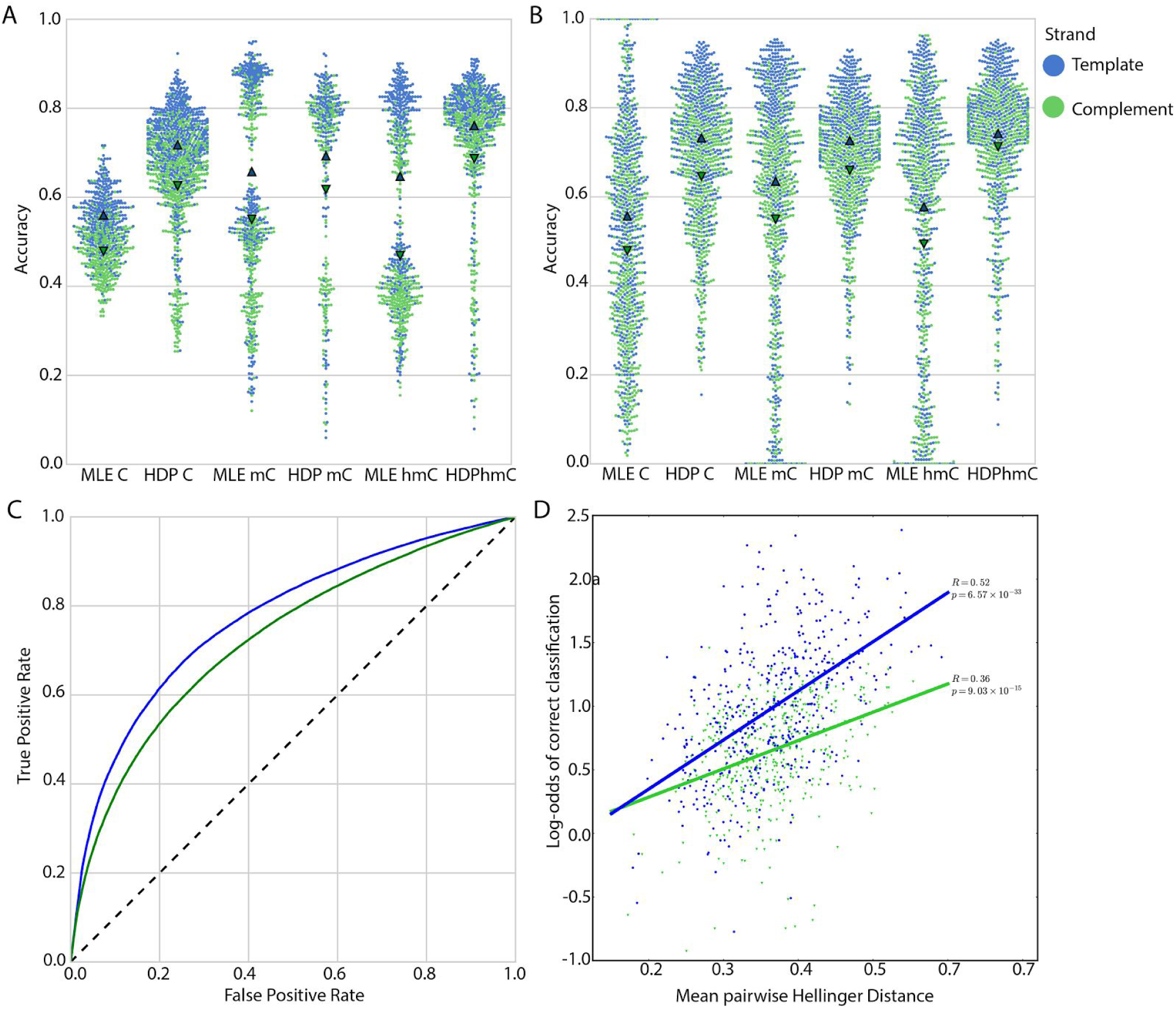
Overview of models and chemical structures. A. Chemical structures of three cytosine variants. B. Architecture of hidden Markov model used in this study. The match state ‘M’ emits an event-6-mer pair and proceeds along the reference, Insert-Y ‘I_y_’ emits a pair but stays in place, and Insert-X ‘I_x_’ proceeds along the reference but does not emit a pair. C and D. Two-level (C) and three-level (D) hierarchical Dirichlet process shown in graphical form. Circles represent random variables. The base distribution ‘H’ is a normal inverse-gamma distribution for both models. The Dirichlet processes ‘G_0_’, ‘G_σn_’, and ‘G_σni_’ are parameterized by their parent distribution and shared concentration parameters ‘γ_B_’, γ_M_’, and γ_L_’. The factors ‘θ_ji_’ specify the parameters of the normal distribution mixture component that generates datum ‘x_ji_’.

**Figure 4.**
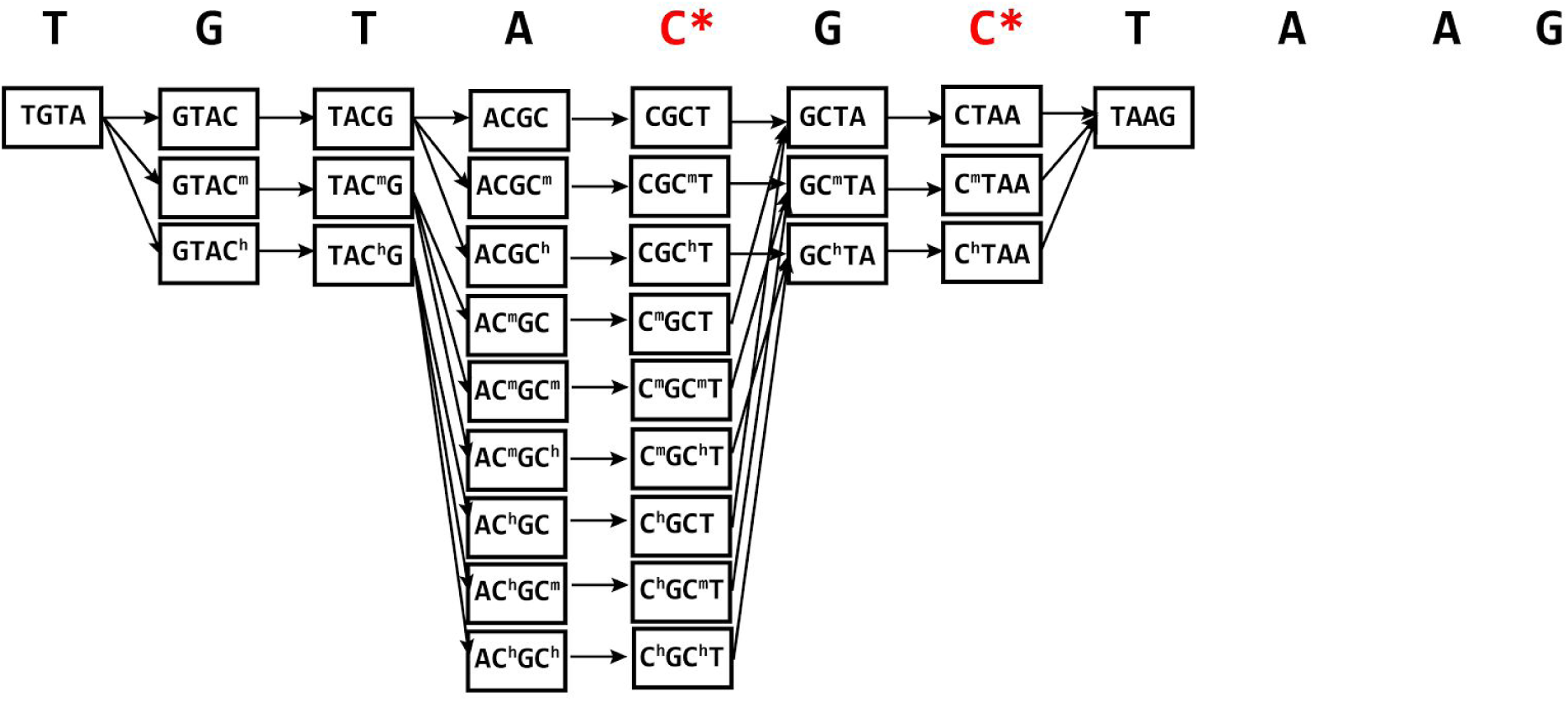
Illustration of variable order hidden Markov model meta-structure. The HMM’s meta-structure ties the possible match states so that ambiguous positions are labeled consistently within a path. Each cell contains the three states shown in Figure 1B, and transitions span between cells. An example reference sequence is shown at the top. C* represents a potentially methylated cytosine. The structure expands around the C* base to accommodate for all possible methylation states. The states are drawn as 4-mers for simplicity, but the model is implemented with 6-mers.

This model allows for simultaneous reference alignment and probabilistic calling of DNA modifications. We show that it can accurately distinguish DNA modifications using synthetic DNA substrates containing homogeneously methylated, hydroxymethylated, or unmethylated cytosine residues.

## Results

### Methylation variant calling

We sequenced synthetic DNA strands containing entirely either cytosine, 5-methylcytosine, or 5-hydroxymethylcytosine on the MinION using standard preparation protocol (see Methods for details). During sequencing, the MinION records ionic current in real time at 3 kHz and then divides it into “events” that correspond to a single nucleotide step of the DNA molecule passing through the nanopore. The current software (and our method) models each event as being due to six nucleotide segments of DNA, which we refer to as 6-mers. A hairpin is ligated to the end of the DNA duplex during sample preparation so that both the template and complement strands are sequenced. We separately align the events from the template and complement strands to a reference sequence with our model and marginalize over the HMM’s states to obtain the posterior probability on the methylation status of a given cytosine. We then call the variant as the methylation status with the highest marginal probability. We performed three-way classification experiments between all three cytosine variants and two-way classification experiments between only cytosine and 5-methylcytosine. The error rates for each read (across cytosines) and each cytosine (across reads) are summarized below.

### Methylation calling error rate

The mean and median per-read accuracy using the best performing HMM-HDP model were 74% and 80% respectively for the template reads and 67% and 76% for the complement reads. The distribution of per-read accuracies is shown in Figure 2A. These results represent a significant improvement over the 33% accuracy that would be expected by chance. They are also significantly better than the results of the HMM with the emissions modeled by normal distributions, which achieved mean and median accuracy of 58% and 62%, respectively, for the template reads and 47% and 50% for the complement reads. When the HMM-HDP classifies between only cytosine and 5-methylcytosine, the mean and median accuracy increase to 83% and 85% respectively for template reads and 78% and 84% for complement reads (Table 1).

**Figure 2.**
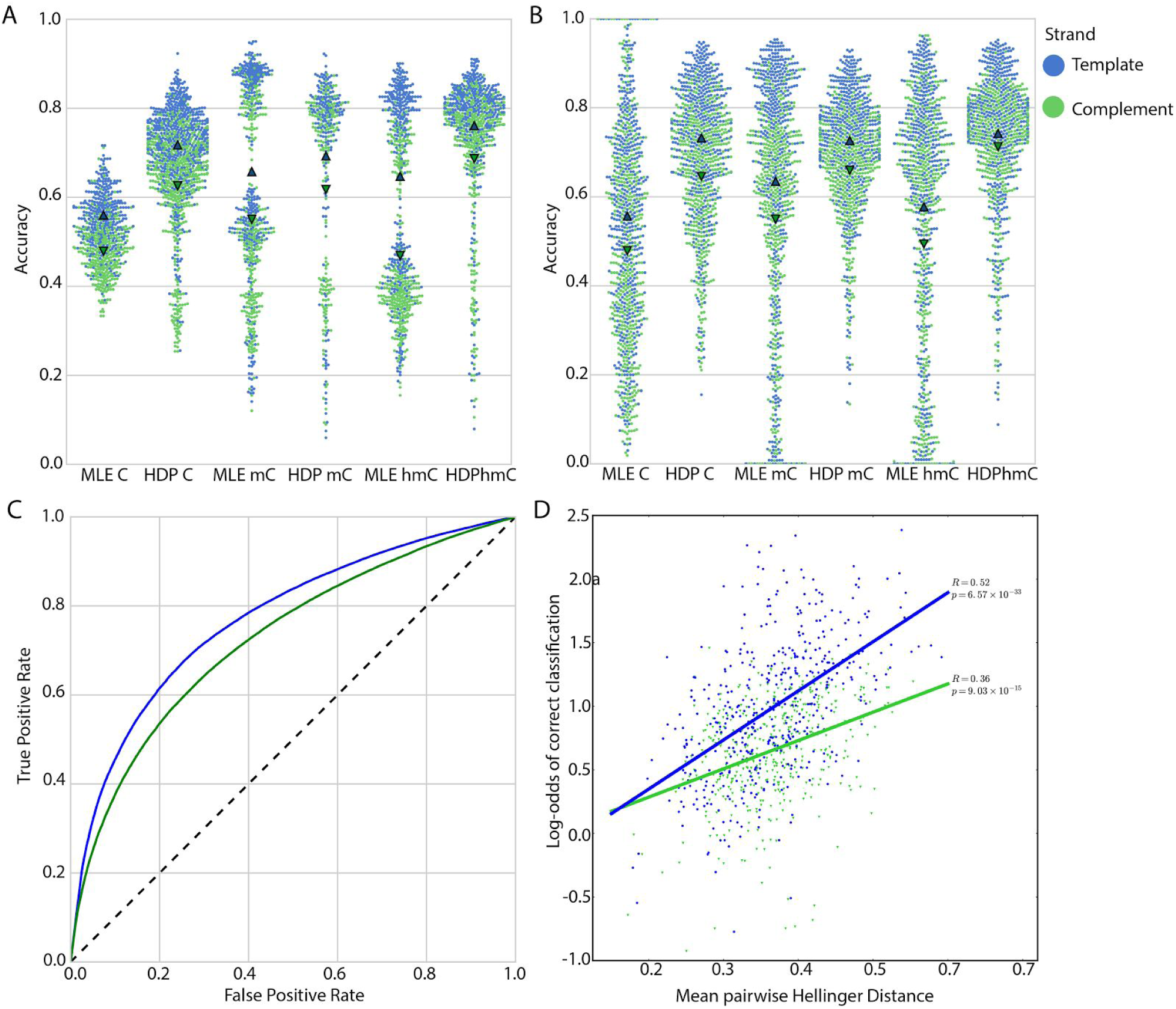
Distributions of accuracy by sequence context and read, ROC plot, and correlation between Hellinger distance and accuracy. A and B. The accuracy distribution by read (A) and by context (B) is shown for the MLE emission distributions and the ‘multiset’ HDP model. The triangles represent the mean of the distribution. C. Receiver operating characteristic plot showing the performance of the classifier across posterior match probabilities. D. Scatter plot shows the correlation between log-odds of correct classification and the mean pairwise Hellinger distance between the methylation statuses of the 6-mer distributions overlapping a cytosine.

**Table 1.** Comparison of different methods and HDP topologies. MLE is the maximum likelihood estimate, described in the text. ‘Two-level’ is an HDP model with no subgroupings of 6-mers, ‘Multiset’, ‘Composition’, ‘MiddleNucleotides’, and ‘GroupMultiset’ are three-level HDP models described in the results. Three-way classification was performed between cytosine, 5-methylcytosine, and 5-hydroxymethylcytosine. Two-way classifications were between cytosine and 5-methylcytosine.

|  |  |  |  |  |
| --- | --- | --- | --- | --- |
|  |  | <u>Three-Way</u><br><u>Accuracy</u> |  |  |
| <u>Model</u> | <u>Mean Accuracy</u><br><u>(read)</u> | <u>Median Accuracy</u><br><u>(read)</u> | <u>Mean Accuracy</u><br><u>(site)</u> | <u>Median Accuracy</u><br><u>(site)</u> |
| <u>MLE</u> | 62% / 50% | 58% / 47% | 59% / 51% | 66% / 57% |
| <u>singlelevel</u> | 74% / 66 % | 79% / 72% | 73% / 63% | 76% / 69% |
| <u>multiset</u> | 74% / 67% | 80% / 76% | 73% / 67% | 76% / 70% |
| <u>composition</u> | 73% / 66% | 78% / 71% | 73% / 66% | 76% / 69% |
| <u>middleNts</u> | 71% / 63% | 76% / 69% | 72% / 64% | 75% / 67% |
| <u>GroupMultiset</u> | 73% / 65% | 78% / 71% | 72% / 66% | 75% / 69% |
|  |  | <u>Two-Way</u><br><u>Accuracy</u> |  |  |
| <u>Model</u> | <u>Mean Accuracy</u><br><u>(read)</u> | <u>Median Accuracy</u><br><u>(read)</u> | <u>Mean Accuracy</u><br><u>(site)</u> | <u>Median Accuracy</u><br><u>(site)</u> |
| <u>singlelevel</u> | 83% / 78% | 86.5% / 84.5% | 82% / 77% | 83% / 78% |
| <u>multiset</u> | 83% / 78% | 86.5% / 84.5% | 82% / 77% | 83% / 78% |

The accuracy varied substantially between different sites on the DNA substrate (Figure 2B). Averaged across reads, the best-performing three-way model classified cytosines at accuracies ranging from 16% to 95% with median accuracy of 76% for template reads and 70% for the complement reads (Table 1). The highest accuracy was achieved in a two-way classification at 98% on template reads, with a median accuracy of 82%. The variability in accuracies agrees with previous research that showed that sequence context affects methylation-calling error rate^13,16^. Figure 2C shows the classifier’s tradeoff between false positive and false negative rate by site across thresholds on the posterior probability.

It is likely that some of the difficulty in classifying certain sites results from 6-mer ionic current distributions that vary only slightly between the methylation states. We observed a statistically significant correlation between the mean pairwise Hellinger distance between the distributions of the methylation states of the 6-mers overlapping a site and its classification accuracy: Pearson correlation 0.52 (p = 6.6E-31) on the template strand and 0.36 (p = 9.0E-15) on the complement strand (Figure 2D).

### The hierarchical Dirichlet process more realistically models ionic current distributions

Figure 3 compares the current signal distributions of three representative 6-mers from the HDP, the maximum likelihood estimate (MLE) normal distribution, and a kernel density estimate. Qualitatively, compared to MLE, the HDP posterior densities reflect the nuance of the 6-mer distributions more realistically. As a nonparametric method, the HDP can approximate any empirical distribution with sufficient data. The statistical shrinkage between the distribution estimates also tends to smooth away small-scale irregularities that can be observed in the kernel density estimate.

**Figure 3.**
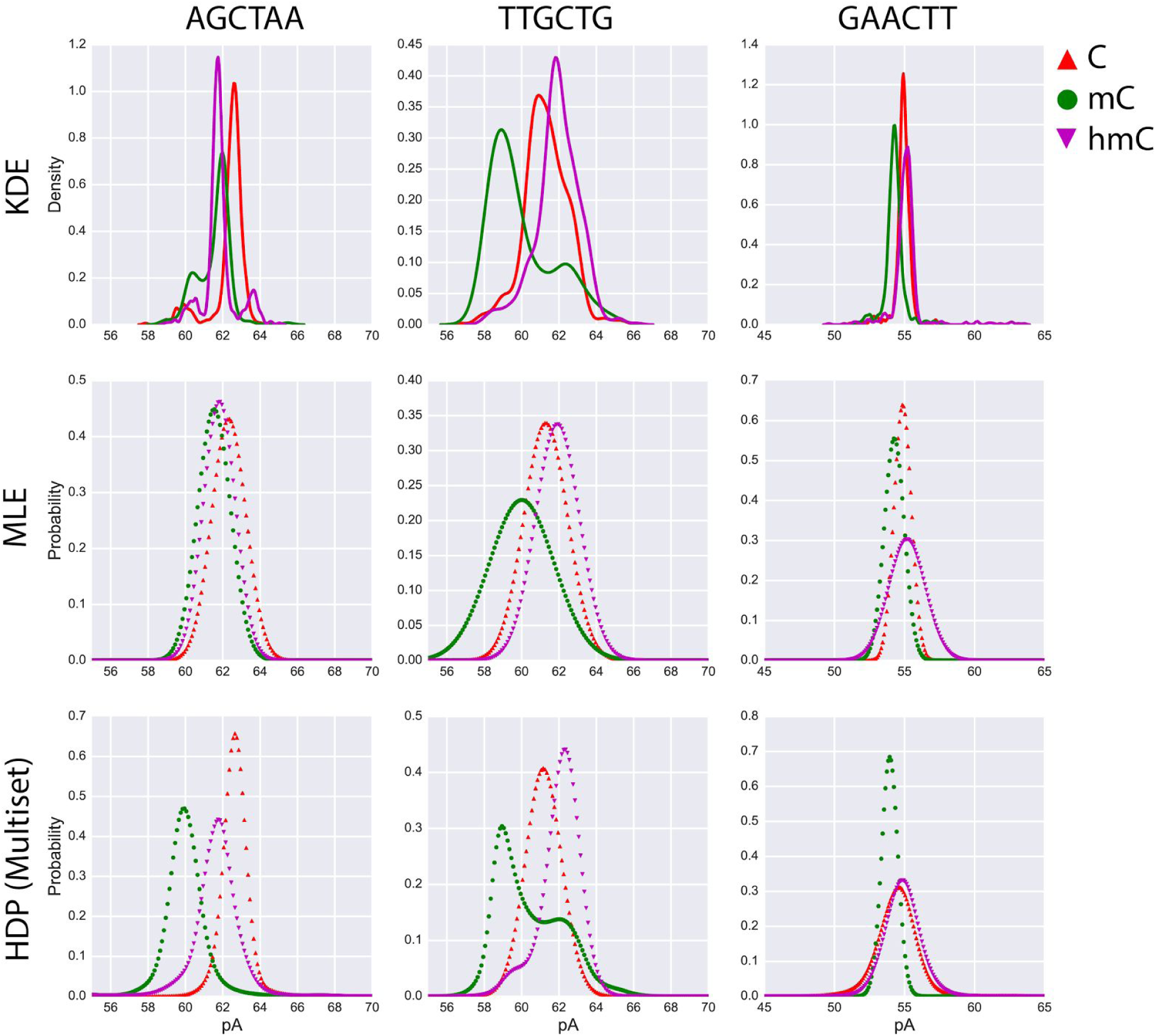
Probability distributions for three representative 6-mers by multiple methods. The first row shows the kernel density estimate (KDE) based on the preliminary alignments described in the text. The middle row shows the probability density function maximum likelihood estimated normal distribution (MLE). The bottom row shows probability density functions from the ‘multiset’ hierarchical Dirichlet process (HDP). All data shown are from template reads.

### Comparison of different HDP topologies

The HDP boosts its statistical strength by sharing information between the set of distributions it estimates. In effect, this encourages them to be more similar to each other than if they were modeled independently. The HDP model also has the possibility of encouraging a greater degree of similarity between pre-specified subgroups of distributions (see Methods for details). This can increase statistical strength further, assuming that the subgroups reflect clusters of similarity in the true distributions. Since the biophysical relationship between each given 6-mer sequence and the observed ionic current distribution is poorly understood, we empirically tested whether certain subgroupings would be informative in this manner.

We tested five HDP models with different subgroupings of 6-mers. The two-level HDP does not separate them into any subgroups (Figure 1C), whereas the rest of the models group 6-mers by features of their 6-mer sequence (Figure 1D). The “Multiset” HDP groups 6-mers by their nucleotide content without regard for the order. “Composition” groups 6-mers by how many purines and pyrimidines they contain. “MiddleNucleotides” groups 6-mers based on the center two bases in the 6-mer. Finally, “GroupMultiset” groups the 6-mers by their nucleotide content without regard for their order or their methylation status. We used methylation-calling accuracy to assess the performance these structures. The best performing model was the “Multiset” model (Table 1). However, it was a small gain in accuracy over the simpler ungrouped model.

## Discussion

To date, few sequencing technologies have been able to directly sequence modified bases alongside canonical nucleotides. ONT’s standard statistical model for the MinION also does not distinguish 6-mers according to methylation. Our results show that it is possible to expand the nucleotide alphabet to include 5-mC, and 5-hmC using a hybrid statistical model composed of a pair HMM and an HDP mixture of normal distributions.

We demonstrate that high-throughput nanopore sensing can successfully discriminate between cytosine, 5-methylcytosine, and 5-hydroxymethylcytosine. Using MinION signal data, we achieved three-way and two-way classification accuracy up to 95% and 98%, respectively of single cytosines and median accuracies of 80% and 85% by read. The classification accuracy varies between sequence contexts: some modified cytosines are reliably captured while others are not discernable. We only classified cytosine variants based on one strand, however, and in an application where there is symmetric methylation the context on the reverse complement strand may be more accurately classified. Rereading uncopied DNA may also improve the accuracy.

We anticipate numerous biological applications for this technology. In particular we expect the combination of long reads and detection of multiple base modifications to be widely useful. For instance, it could be applied to studying genomic methylation and haplotype phasing. Since no extra sample preparation is necessary, this information is available “for free” in any sequencing experiment. With appropriate training data, our methodology could be easily generalized to detect additional nucleotides and different base modifications as well. As nanopore sequencing evolves, the accuracies for detecting base modifications will improve further, opening this technology to diagnostics and other clinical applications.

## Methods

### Creating a controlled set of C, 5-mC, and 5-hmC sequences

We used 897 bp synthetic DNA strands from ZYMO Research (Catalog # D5405) that contain entirely C, 5-mC, or 5-hmC bases. Apart from the cytosines, the strands have identical sequences. We performed sequencing experiments (using SQK-MAP006 kits) with four MinION flow cells: one for each of the three substrates, and one where all the substrates were with barcoded with uniquely identifying sequences (using an ONT kit) and run together on one flow cell. All models were trained on the reads run in separate flow cells. The bar-coded reads served as our test dataset. This experimental design maximized the amount of training data while controlling for batch effects between MinION runs. Sequence data were processed using Metrichor (versions 1.15.0 and 1.19.0), and only ‘pass’ 2D reads were used for downstream analysis.

### Mapping of Reads and Event Alignment

We align ionic current events to the reference sequence in a two-step process. First we generate a guide alignment between nucleotide sequences, which we then use to guide a second alignment of events to the reference. To generate the guide alignments, we used a concatenated sequence from Metrichor’s ‘2D alignment’, which allows for each base in the MinION nucleotide sequence to be mapped to an event in the template and complement event sequence. We then generated a guide alignment of the nucleotide sequence to the reference with BWA-MEM in ont2d mode^17^. Runs of consecutive matches in this guide alignment serve as anchors for the event-to-reference alignment using the banded alignment scheme described by Paten et al.^18^. The anchors are mapped back to events in the event sequence, and the events are then realigned to the reference using the HMM described below, constrained by the anchors.

### Structure of variable-order hidden Markov model

Our HMM is structured to allow alignment of multiple different bases at a given position in the reference sequence. In this study, we allow for any cytosine variant to be aligned to a given cytosine residue. The fact that each event corresponds to six positions in the reference means that more than one event reports on a single ambiguous position. Accordingly, the HMM must be constrained so that two nearby match states cannot label a reference cytosine’s methylation inconsistently. To accommodate this, we implemented our HMM in a variable-order meta-structure that allows for multiple paths over a reference 6-mer depending on the number of methylation possibilities (i.e. the number of cytosine options raised to the power of the number of ambiguous positions in the 6-mer). The dynamic programming matrix has high-dimensional cells to accommodate these paths. We restrict the recursion by only allowing transitions if the bases at positions 2-6 in the first 6-mer are identical to the bases at positions 1-5 in the second 6-mer (Figure 5). The joint probability for the event sequence and the reference is calculated with the forward-backward algorithm, and the likelihood of methylation at each cytosine is calculated by marginalizing over the HMM’s states.

### Hierarchical Dirichlet process mixture model

The HDP mixture is a statistical model in which a collection of mixture distributions (here corresponding to the signal emission distributions for the 46,656 different 6-mers in the expanded alphabet) are composed of a countably infinite set of shared mixture components. The weights of the components in each mixture distribution are determined according to a separate Dirichlet process on the shared collection of components^19^. In addition, the mixture components themselves are distributed according to a Dirichlet process that draws components from a base distribution. In our model, the base distribution is the normal-inverse gamma distribution, which is a conjugate prior to the normal distribution (that is, to the mixture components).

Sharing mixture components statistically shrinks our estimates of the current distributions toward each other. This boosts statistical strength since each distribution can share the information learned by the others. We also have the option of adding a further layer of Dirichlet processes between the Dirichlet process that generates the distribution over shared components and the Dirichlet processes that generate the 6-mer distributions. After doing so, the Dirichlet processes are arranged in a tree structure (Figure 1D). This encourages a greater degree of shrinkage within each subtree. We experimented with several topologies for this tree, each representing a different grouping of 6-mers based on their sequence composition (see Results for descriptions of the groupings).

### Generating preliminary alignments without consideration for methylation status

ONT provides a lookup table of parametric distributions that they use characterizes the current distributions of the 4096 canonical base 6-mers. We take advantage of this table to heuristically initialize the emission distributions in our HMM over the expanded alphabet. To do so, we generate a preliminary alignment using the table and then infer the methylation status of the events based on their flow-cell (as mentioned above, the substrates within a flow cell only contain one kind of methylation). We can then use high probability matches from this alignment to train the emission distributions of the HMM.

To generate preliminary alignments we used the ONT table to calculate the probability an event being due to a particular 6-mer in the Match and Insert-Y states of the HMM. The event’s mean current and fluctuation in the mean (noise) are modeled as normal distributions. We assume independence of the mean and noise variables, so the conditional probability of an event for a given 6-mer is just the product of the mean and noise marginal probabilities. The Insert-X state is silent and therefore does not have an emission probability.

### Supervised training of 6-mer distributions

We train the HMM with a variant of the Baum-Welch procedure. First, we heuristically initialize the emission distributions by training them on aligned events above a probability threshold (0.9) from the preliminary alignment described above. In the control experiments using normal distributions, this simply entails calculating the maximum likelihood normal distribution for each 6-mer. For the HDP-HMM, we estimate the posterior mean density for each 6-mer’s distribution using a Markov chain Monte Carlo (MCMC) algorithm. In both cases, we only estimate distributions for the event mean current following the preliminary alignment (a separate neural net experiment suggested that the event noise did not add to classification accuracy; Supplementary Methods). We then produce new alignments and re-estimate the emission distributions from high confidence assignments as in the initialization. At this step, we also re-estimate the HMM’s transition probabilities independently. This process is iterated until the model’s variant calling accuracy stops improving.

The MCMC algorithm we use for the HDP is the Chinese Restaurant Franchise Algorithm (Teh, et al. 2006), a Gibbs sampler for HDP mixture models. We discard the first 900,000 samples as burn-in (30-times the total number of assignment data points) and collect 10,000 samples, thinning sampling iterations by 100. Whenever we record samples from the Markov chain, we evaluate the posterior predictive distribution for each 6-mer at a grid of 1200 evenly spaced points in the interval between 30 pA and 90 pA. After sampling, we compute our estimate of the posterior mean density as the mean of the sampled densities at each grid point. Subsequently, we interpolate within the grid using natural cubic splines.

## Supplementary Information for ‘Cytosine Variant Calling with High-throughput Nanopore Sequencing’

**Supplementary Methods**

**1 Preparing event and nucleotide sequences**

**Making a read sequence from the 2D alignment table**

The current MinION DNA sequencing library preparation protocol involves ligating a DNA hairpin to the distal end of the substrate to be sequenced. This effectively makes the DNA duplex one long strand of nucleic acid polymer. The sequencer then proceeds to sequence both strands of the DNA duplex. When these two “1D” reads are basecalled there is an event attributed to each nucleotide in the read. The two 1D reads are assembled *in silio* into a “2D” read. During the assembly process some bases are inferred by the algorithm and do not have events attributed to them. We need every nucleotide in the sequence to correspond to an event because we use BWA-MEM to map the nucleotide reads to the reference sequence and use runs of consecutive matches to constrain the dynamic programming (see Banding section below). Obviously, the inferred bases cause problems when we try to map bases to events. The “2D Alignment” table, however, does attribute each nucleotide position to an event. Therefore, we use the list of 6-mers in the 2D alignment table to construct our nucleotide sequence, which is then used for all downstream analysis.

**Descaling events**

The software that processes the raw MinION signal into events also produces three read-specific parameters that describe how the distributions of the signal from that read deviate from Oxford Nanopore’s standard statistics: “scale”, “shift”, and “variance”. Scale indicates how spread out the distributions of each 6-mer are from each other, shift indicates the location of the entire set of distributions, and variance indicates how flat or peaked each distribution is (although it is actually a standard deviation, not a variance). Without these parameters, the standard statistics describe each event mean *X* as a normal distribution with a given mean and variance:

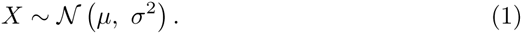

The deviation parameters indicate that, from this particular read, *X* actually follows

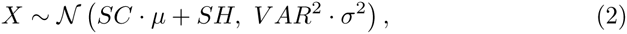

where *SC*, *SH*, and *V AR* are the scale, shift, and variance parameters respectively. Rather than relying on the standard statistics, we learn 6-mer distributions with a hierarchical Dirichlet process (See below). The learning algorithm we use does not permit us to reparametrize the signal distributions for each read. Instead, we reverse the process. Rather than transforming the *distribution*, we transform the *signal*. In particular, we use a change of variable transformation that would transform (2) into (1) so that all of the events can be assumed to use the same coordinates. It is straightforward to verify that this transformation is

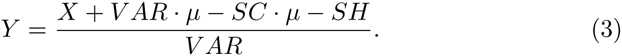

We then use our model learn the density of *Y* and backtrack to obtain a read-specific density for *X* (accounting for the Jacobian of the transformation):

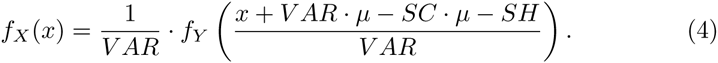

**2 Modeling Ionic Current Sequences**

During strand sequencing multiple nucleotides occupy the nanopore at one time and thus multiple nucleotides contribute to the ionic current. In this study, we model the ionic current as being due to six nucleotides (6-mers) but other models with different length kmers could been used. The motor enzyme moves the DNA one nucleotide at a time through the nanopore so five of the nucleotides remain and one changes, resulting in a new 6-mer in the nanopore. In this section we describe a pair hidden Markov model (HMM) for aligning ionic current events (event sequence) to nucleotides (reference sequence). We then describe how this model is expanded to allow for alignment to multiple variants at a positions in the reference sequence.

**2.1 Pair Hidden Markov Model**

A pair-HMM with the structure shown in Figure 1A was implemented, the code can be found at http://https://github.com/ArtRand/signalAlign. This model calculates the probability of an alignment *π* given a sequence of events *E* = {*e*_1_…*e*_n_}, a sequence of nucleotides, *S*, divided into nucleotide 6-mers *S* = {*k*_1_…*k*_n_}, and the model *Θ*; *P*(*π*|*E*, *S*, *Θ*). The three states; match *M* for matching one event with one nucleotide kmer, *I*_*x*_ for pairing a nucleotide kmer with a gap, and *I*_*y*_ for pairing an event with a gap. The transition probabilities are initialized to naive estimates for the stride, skip, and stay probabilities that correspond to the enzyme advancing exactly one nucleotide, advancing more than one nucleotide, and not advancing, respectively. In one version of the model, the emissions for the *M* state and *I*_*y*_ are the product of the probability of the ionic current mean and ionic current noise. We assume independence of the mean and noise variables, so the conditional probability of an event for a given 6-mer is given by,

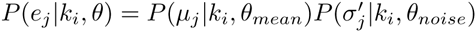

The mean and noise are modeled as a normal distributions *μ*_*i*_ ~ *Ν*(*μ*_*i*_, *σ*_*i*_) and 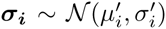, where *μ*_*i*_, *σ*_*i*_, 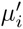, and 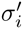 are given by the supplied table from Oxford Nanopore Technologies. For alignments using the maximum likelihood estimate (MLE) (see methods in text for details of MLE), we only update the *μ*_*i*_, *σ*_*i*_ parameters. The *I*_*x*_ is silent and does not emit. In another version of the model we modeled the ionic current distributions as a hierarchical Dirichlet process mixture of normal distributions (details below). In this case we use the posterior mean density as the emission probability of the *M*, and *I*_*y*_ states instead of the probability density function for a normal distribution.

**2.2 Variable-Order Hidden Markov Model**

We model each event as reporting on six nucleotides in the reference sequence. When we allow for multiple variants in the reference sequence (multiple cytosine variants) we would like to compute over all possibilities in a way that the probabilities for a given 6-mer are tied with only 6-mers that share that particular variant pattern. As can be seen visually in Figure 5, when a position is allowed to be variable (C* bases) the number of paths expands to accommodate the number of variable positions. Given 6-mer *k*_*i*_ that contains *η* variable bases within the set *C* = {*A*, *C*, *G*, *T*, *C*^*m*^, *C*^*h*^} the number of paths, *l*, is simply *l* = *η*^|*C*|^. The dynamic programming matrix is changed such that every cell has *l* dimensions, which is precomputed based on the reference. Then we perform the forward-backward algorithm through the matrix except that we don’t want to sum over all paths, *Π*, only paths that represent legal moves, *π* ⊆ *Π*. A move is defined as legal if bases 2-6 of the previous path’s 6-mer are the same as bases 1-5 of the current path’s 6-mer. For example, assume 5m-C is represented at E and 5hm-C is O, the move between 6-mers `AGEOAT` and `GEOATA` would be legal, but the move between `AGEOAT` and `GEEATA` would not. Moves from the start state and to the end state are legal regardless of the 6-mer. With this framework we can calculate the total probability of an event sequence, the reference sequence containing variable positions, and the model by:

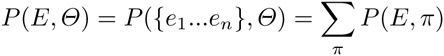

Next we can calculate the posterior probability that 6-mer *x*_*i*_ is aligned to event *e*_*j*_ (noted as *x*_*i*_ ⋄ *e*_*j*_) as

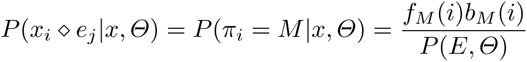

Where *ƒ*_*M*_ and *b*_*M*_ are the forward and backward variables respectively.

**2.3 Banded alignment**

We use a banded alignment heuristic to increase the speed and memory requirement of our algorithm. The banding procedure is described in detail in [1], in this section we briefly describe the procedure and the settings used. The nucleotide sequence derived from the 2D alignment table (described above) is aligned to the reference sequence using BWA-MEM using the-ont2d flag for aligning nanopore reads. We refer to this as the guide-alignment. From the guide alignment runs of un-gapped matches are used as constraints in the edit graph around which we compute our dynamic programming. To prevent any edge effects, the constraints are trimmed at either end by 14 nucleotide pairs. We then expand around the constraints by 50 anti-diagonal cells in the edit graph. To increase the efficiency of the algorithm, we break the alignment into fragments. Alignment bands computed between the centers of the constraints where the quality of the guide alignment should be highest. This allows our higher order HMM to have a smaller memory footprint as well as constraining the posterior probability distribution to higher likelihood matches.

**2.4 Hierarchical Dirichlet Process Mixture Model for Ionic Current Distributions**

We model the distribution of ionic currents across the 45,656 different 6-mers as a hierarchical Dirichlet process (HDP) mixture of normal distributions. In this model, each current distribution is composed of a countably infinite collection of Gaussian mixture components that are shared between the 6-mers. More precisely, all of the distributions draw mixture components according to a Dirichlet process over the same discrete distribution over a countably infinite collection of “atoms”, each of which consists of the parameters for a normal distribution. In addition, this discrete distribution is itself generated according a Dirichlet process on the normal-inverse-gamma distribution (a conjugate prior to a normal distribution).

The most intuitive interpretation of the Dirichlet process for this setting is the “stick-breaking procedure”. In this construction, we draw a countably infinite number of atoms from the normal-inverse-gamma distribution and then form a new distribution over these atoms by assigning them weights *w*_k_, *k* = 1,…,∞ sequentially:

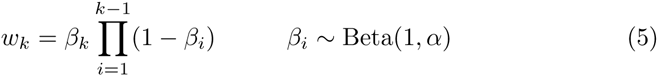

where *α* is a hyper-parameter referred to as the “concentration”. Note that both the value of the atoms and their weights are random variables. To generate each of the current distributions, the HDP draws a countably infinite set of mixture components according to a Dirichlet process on this new distribution (treating the weights as probability masses). The weights that this process assigns to the atoms are the weights on the components in the mixture distribution.

Our motivation for using an HDP to model signal distributions is that it shrinks the set of distributions it learns towards each other, which increases robustness, while retaining the flexibility to approximate any arbitrary distribution given sufficient data. Both of these feature are important, since we expect that the current distributions may have complex shapes and we must estimate a large number of them. The shrinkage is a result of the fact that all of the distributions share the same mixture components: each distribution can share the information learned by the others. In addition, this allows us to calculate informed prior distributions even for 6-mers that have not been positively identified in the training data, a feature that will be useful for expanding the scope of modifications that the model can detect.

**Determining appropriate hyperparameters**

The HDP mixture has two sets of hyperparameters. First, we must choose the concentration parameters. Rather than choosing fixed values, we use the methods of [2] and [3] to place an exponential distribution prior with expected value 1 on the concentration parameters. We also assume that each level in the hierarchy of Dirichlet processes shares one concentration parameter. In other versions of the model, we experimented with fixed concentration parameters and with exponential priors with larger expected value, but this version performed the best.

We also must choose the parameters of the normal-inverse-gamma distribution that generates the atoms for the base Dirichlet process. To do so, we leverage information provided by Oxford Nanopore Technologies. MinION data come with a table of normal distributions that describe current signal generated by the 4096 six-nucleotide 6-mers composed of the four canonical bases. We expect the set of distributions that HDP learns to be broadly similar to these current distributions in terms of location and scale. Accordingly, we chose a base distribution for the HDP’s mixture components to have a high likelihood of generating the normal distributions described in the table. In particular, we found the maximum likelihood normal-inverse-gamma distribution treating the distributions in the table as previously observed mixture components according to the following derivation. For the sake of cleaner equations, we solve the equivalent problem of finding the maximum likelihood estimates for the parameters of a normal-gamma distribution.

Given *N* independent observations of mean-precision tuples (*μ*_*i*_,*τ*_*i*_), we have likelihood function

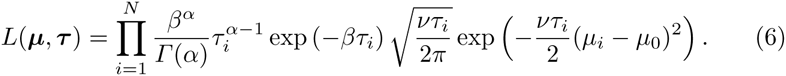

Differentiating the log-likelihood, the maximum likelihood estimates then satisfy

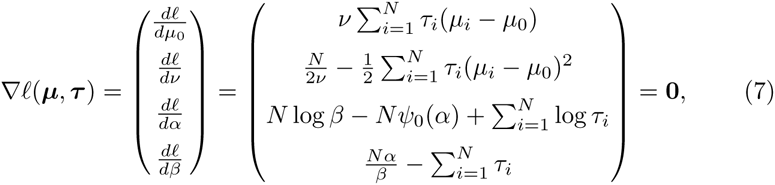

where *ψ*_0_ is the digamma function. This immediately yields closed form solutions for the parameters *μ*_0_ and *v*:

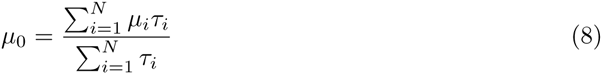

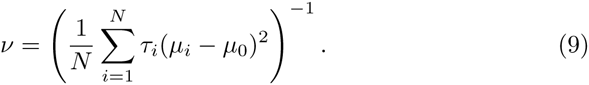

Further, substituting

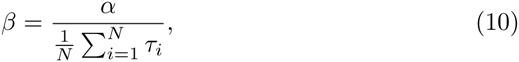

we obtain

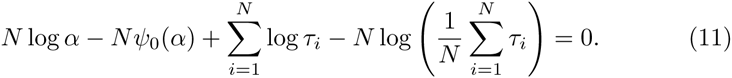

This formulation permits a Newton’s method approximation of *α*. In particular, starting from arbitrary positive *α*_0_ we iterate the following until convergence:

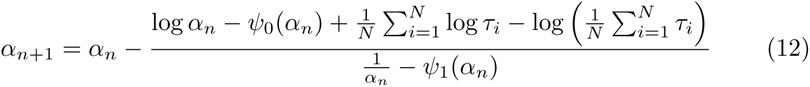

where *ψ*_1_ is the trigamma function. The result then also gives a solution for *β* through Equation (10).

**Grouping 6-mers with different HDP topologies**

In the original HDP described by Teh, et al. (2006), all of the mixture distributions draw mixture components from the same distribution over atoms, which is generated by a Dirichlet process on the base distribution. However, there is a relatively simple extension of this model in which the mixture distributions are split into prespecified groups, and it is these groups that share a distribution of over atoms. This is accomplished by adding an additional layer of Dirichlet processes between the one that generates the initial distribution over atoms and the ones that generate the mixture distributions. To conceptualize this, it helps to think of the HDP as a collection of Dirichlet processes arranged in a tree structure (with the Dirichlet process over the base distribution at the root). With this framing, the generalization amounts to having a tree with a depth of two instead of one.

All of the mixture distributions still share the same collection of atoms, since all of the atoms in the middle layer of Dirichlet processes are drawn from the original root Dirichlet process. However, the weights of the atoms are reassigned according to a new stick-breaking procedure (See Equation (5)) in each of the middle-level Dirichlet processes. The effect is that the shrinkage between the distribution estimates is greater within a subtree than between subtrees. Since we have control over the topology of the tree, this serves an extra “knob” that can be used to increase statistical strength. However, the grouping of mixture distributions into subtrees presumably must reflect clusters of similarity in the true distributions in order to accomplish this goal.

The biophysics of how a 6-mer of DNA translates into a current distribution is poorly understood. Accordingly, we took an empirical approach to determining what topologies for the tree of Dirichlet processes would be meaningful. We came up with several ways to partition 6-mers based on their sequence composition and tested the performance of each one. The groupings were as follows:

1. *No groups*: this model has no middle layer of Dirichlet processes.
2. *Groups of 6-mers containing the same number of purines and pyrimidines*: this corresponds to a hypothesis that steric bulk is a strong determinant of the current distribution.
3. *Groups of 6-mers that shared the same two middle nucleotides*: this corresponds to a hypothesis that the nucleotides that are passing through the most constricted portion of the nanopore have the most influence.
4. *Groups of 6-mers that contain the same nucleotides, irrespective of their order*: this corresponds to a hypothesis that the position of nucleotide in the 6-mer is less important than which nucleotide it is, similar to resistors in series.
5. *Groups of 6-mers that contain the same nucleotides, irrespective of both their order and their modification status*: this corresponds to the same hypothesis as the previous grouping and also a hypothesis that the modification status has relatively little effect on the current distribution.

**2.5 Training the HMM Transitions**

We trained the HMM transitions separately from the emissions using a batch expectation-maximization procedure usually referred to as the Baum-Welch algorithm. For a detailed description of the algorithm, see Durbin et al. 1998. Training was performed in batches of 15,000 nucleotides and iterated 20 times. When experimenting with different emissions models (eg. different HDP topologies), the transition matrices were trained specifically for that model.

**2.6 Training the HMM-HDP model**

An overview of the training method used is described in the methods section of the main text. In this section we provide more details for this procedure. We train the HMM-HDP with a variant of the Baum-Welch procedure. However, this algorithm is sensitive to the initial values of the model’s parameters, so first we leverage Oxford Nanopore’s standard statistical model to heuristically initialize the HMM’s emission distributions. ONT’s model consists a table of parametric distributions that describe the events arising from each of 4096 6-mers composed of the four conventional nucleotides. In particular, they model the event mean current as a normal random variable and the event current variability (which they call “noise”) as an inverse Gaussian random variable. We assume that these are independent so that we can calculate their joint density as the product of the marginal densities.

To obtain our heuristic estimates of the HMM-HDP’s emission distributions, we first generate a preliminary alignment with a simplified HMM. This HMM only has states over the conventional four-nucleotide alphabet. This allows us to 1) use the standard statistical model and 2) use a first-order HMM (since there are no ambiguities between the HMM’s alphabet and the reference alphabet). We use the banded alignment scheme described above to obtain a posterior probabilities that an event was generated by a given 6-mer. Our experimental design allows us to then label the methylation status of the cytosines from these alignments post hoc. After doing so, we extract the aligned events with posterior probabilities of at least 0.9 and use these as training data to learn emission distributions for the HDP.

Once we have a set of aligned events as training data, we estimate the emission distributions using an MCMC method. The specific algorithm is a Gibbs sampler for HDPs called the Chinese Restaurant Franchise Algorithm, which we implemented in original software (available at https://github.com/jeizenga/hdp_mixture). Briefly, it involves integrating out the values of the HDP’s atoms and then sampling the full conditional distributions of latent variables that indicates which mixture component generated each data point (See Teh, et al. (2006) for details). We begin with a burn-in period of 30 times as many sampling iterations as we have data points. After this, we record 10,000 samples, thinning by 100 sampling iterations. Every time we record a sample, we compute the posterior predictive distribution of each 6-mer on a grid that covers the range of MinION current signal. We then estimate the mean posterior density at each of these grid points from the sample and interpolate between them using natural cubic splines. These serve as our distribution estimates.

Our variant of the Baum-Welch procedure functions similarly to the heuristic initialization. First, we align the reads to the reference, except now we use the full variable-order HMM and the emission distributions estimated by the HDP. We then extract the assignments with posterior probabilities of at least 0.9 and use these as training data to obtain mean posterior distributions from the HDP. We then iterate this process a fixed number of times. This differs from the true Baum-Welch procedure since we are using posterior mean estimates rather than maximum a posteriori. However, these values are asymptotically equivalent in unimodal posteriors, so this is probably a reasonable approximation.

**3 Classification of Ionic Current Events with Neural Networks**

We investigated the feasibility of cytosine methylation detection by testing whether or not events aligned to a single cytosine could be classified based on their methylation status. We also used this strategy to evaluate which features are the most discriminatory in classification. Artificial neural networks are non-sparse classifiers well suited to this task, in this section we describe classification of events that have been aligned to the reference sequence using the preliminary alignments generated without consideration for methylation status.

**Data processing for single cytosine motifs**

Subsequences of the reference sequence, motifs, were selected that contain a single cytosine among a run of 11 nucleotides. The MinION reads both strands of the DNA duplex and combines these reads into a ‘2D’, high quality read. The motifs we chose contain guanine bases, however, and the complement would therefore contain a variable number of modified cytosine bases and contribute to the classification accuracy (see methods for substrate description). We therefore classified only one strand per read. In the case of forward-mapped reads this was the template strand, in the case of backward-mapped it was the complement strand. To assess the amount of classification bias due to the stands alone, we classified Null motifs that do not contain any cytosines, Figure S1. These Null motifs should be the same between strands and the classifier should report accuracy close to random chance (33%). Events that aligned to positions in the reference corresponding to a motif were culled. When multiple events were aligned to a position, the one with the highest posterior probability was taken. Thus an ordered set of a maximum of 12 observed event mean current levels (6 on the template and 6 on the complement), six observed ionic current noise levels, and six posterior probabilities, was obtained. The difference from the observed mean current level from the expected current level (from the ONT table) given the 6-mer was taken for each observation. The same was done for the noise levels. The differences in the mean current level (*Δμ*), differences in noise level (*Δσ*), and the posterior probabilities (*P*) became the features input to the classifier. We experimented with four different feature sets; *Δμ* alone, *Δμ* and *Δσ*, *Δμ* and *P*, and all three together.

**Network architecture and training routine**

Classification of the feature vectors was performed using a custom artificial neural network implemented in Theano (Bastien et al. 2012). For the individual motif classification a network with two hidden layers and a final softmax layer was used. The dimensions of this network were 50, 10, 3, with rectified linear unit and hyperbolic tangent nonlinearity activation functions used for the first and second hidden layers respectively. For classification, the class with the highest probability from the softmax layer was chosen. We split the dataset into three groups; 10% of the data was held out for testing after the training procedure. Of the remaining, training data, 50% was used as cross-training (validation) and 50% was used in the optimization. An equal number of feature vectors for each category were used in all data sets. Training of the network was done using mini-batch stochastic gradient descent and an annealing learning rate schedule. We used 5 reads per batch and a dynamic learning rate initialized at 0.1%. With decreases in cross-training batch costs, the learning rate was decreased by 10%, if the cross-training batch cost increased, the learning rate was increased by 5%. We used L1 and L2 regularization of 0.01. Lastly, the data was centered and normalized based on the training data set before starting the routine. The model that had the highest cross-training accuracy during the learning process was used for final evaluation of the test set. We performed the same training routine on the Null sites.

**Classification accuracy is maximized using *Δμ* and posterior probability as features**

The accuracy for the different feature sets for the cytosine motifs is summarized in Table 1, the Null motifs are summarized in Table 2. The highest accuracy was obtained when using the mean and posterior probability at 65% on the template reads and 66% on complement reads. Including *Δσ* did not appear to increase the accuracy of the classifier. We classified events culled from alignments generated with the HMM-HDP and the methylation-naive HMM. Both data sets produced similar error rates when classified with the neural network.

**Table 1.** Classification of single cytosine motifs

| Motif Sequence | Forward/Template<br>$\{\Delta\mu\}$ | $\{\Delta\mu, \Delta\sigma\}$ | $\{\Delta\mu, P\}$ | $\{\Delta\mu, \Delta\sigma, P\}$ |
| --- | --- | --- | --- | --- |
| TTTTGCTGAGT | 78.33 | 86.67 | 83.33 | 85 |
| AAGTTCAAAAT | 40 | 51.67 | 46.67 | 35 |
| AGATGCAGGGG | 70 | 73.33 | 83.33 | 76.67 |
| AAGGGCTGGAT | 66.67 | 75 | 68.33 | 61.67 |
| ATTTGCTGAGG | 65 | 78.33 | 58.33 | 73.33 |
| TGGGGCAAATG | 68.33 | 75 | 68.33 | 83.33 |
| GGAATCAAATT | 40 | 40 | 40 | 40 |
| GTGGACAGGAA | 76.67 | 68.33 | 71.67 | 75 |
| AAATTCTTGAA | 46.67 | 56.67 | 56.67 | 58.33 |
| GAAGACGAAAG | 81.67 | 66.67 | 85 | 76.67 |
| AATGTCATGAT | 53.33 | 61.67 | 68.33 | 73.33 |
| GGTTTCTTAGA | 45 | 43.33 | 58.33 | 51.67 |
| TTTTTCTAAAT | 43.33 | 40 | 60 | 60 |
| Average: | 59.62 | 62.82 | 65.26 | 65.38 |
| Backward/Complement |  |  |  |  |
| TTTTGCTGAGT | 53.33 | 51.67 | 64.44 | 66.67 |
| AAGTTCAAAAT | 41.67 | 43.33 | 62.22 | 60 |
| AGATGCAGGGG | 61.67 | 63.33 | 84.44 | 68.89 |
| AAGGGCTGGAT | 63.33 | 63.33 | 73.33 | 73.33 |
| ATTTGCTGAGG | 51.67 | 70 | 75.56 | 73.33 |
| TGGGGCAAATG | 71.67 | 68.33 | 75.56 | 66.67 |
| GGAATCAAATT | 40 | 41.67 | 40 | 57.78 |
| GTGGACAGGAA | 40 | 40 | 46.67 | 40 |
| AAATTCTTGAA | 35 | 40 | 62.22 | 77.78 |
| GAAGACGAAAG | 73.33 | 55 | 80 | 75.56 |
| AATGTCATGAT | 40 | 40 | 57.78 | 62.22 |
| GGTTTCTTAGA | 45 | 46.67 | 68.89 | 66.67 |
| TTTTTCTAAAT | 46.67 | 48.33 | 68.89 | 64.44 |
| Average: | 51.03 | 51.67 | 66.15 | 65.64 |

**Table 2.**
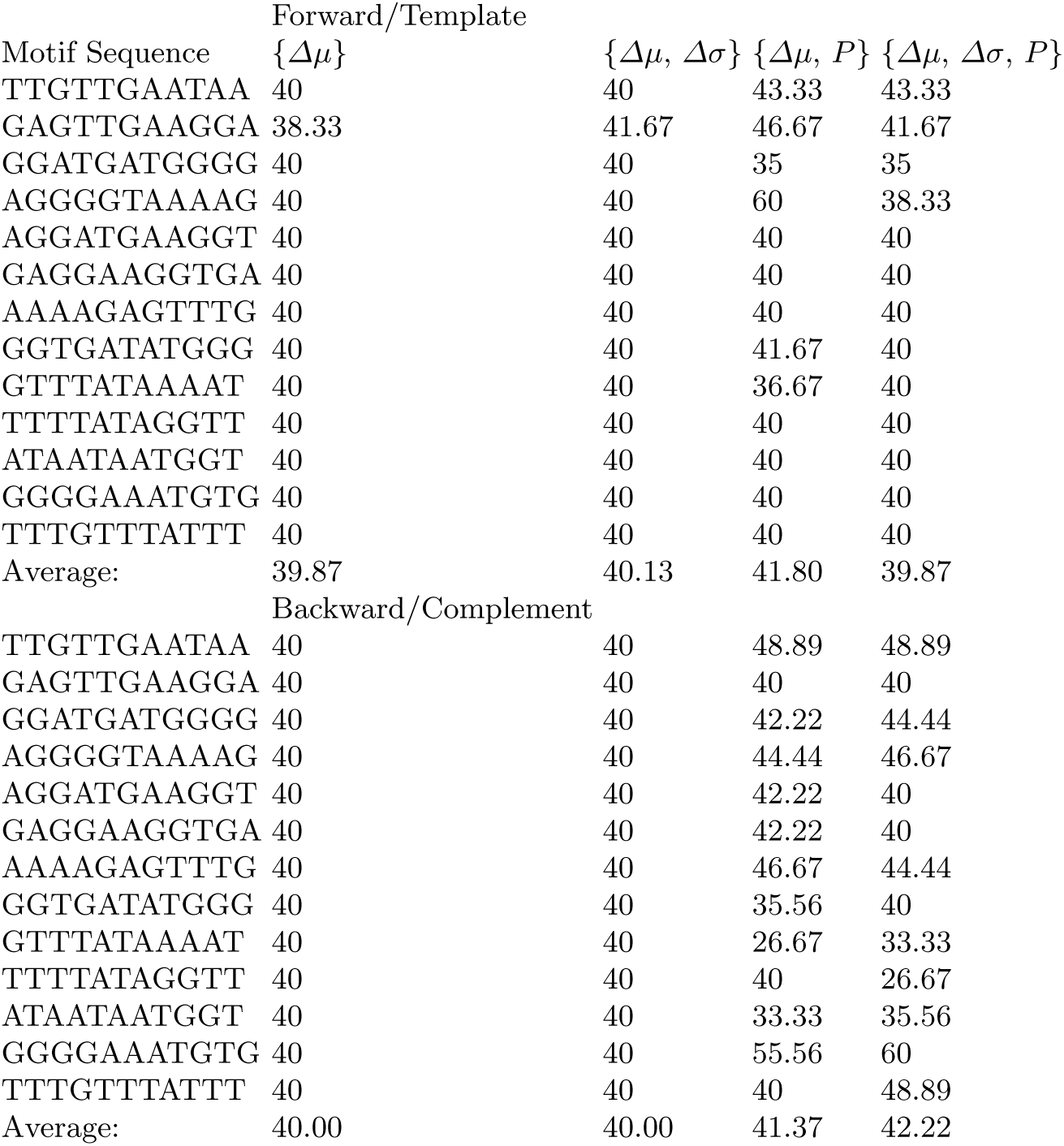
Classification of Null motifs

**4 MinION DNA Sequencing**

We used synthetic DNA substrates from ZYMO Research (Catalog # D5405). These are 897 bp long DNA molecules that are used as 5-mC and 5-hmC DNA standards. We performed three MinION sequencing runs with SQK-MAP006 chemistry kits, using one flow cell per DNA standard (C, 5-mC, and 5-hmC).

Additionally, we also bar-coded these substrates using ONT barcoding kit and mixed them together. We then performed MinION sequencing of this mixture on a single flow cell. Reads corresponding to the three DNA standards were separated using their respective barcode sequences during base-calling. All the data was base-called using Metrichor (versions 1.15.0 and 1.19.0).

For this manuscript, we restricted all downstream analysis to pass 2D reads. We used BWA (in ont2d mode) as the initial alignment tool to obtain read mapping orientations (forward/backward). However, our method is not restricted to 2D reads (or pass/fail categories).

## References

1. Robertson, K. D. DNA methylation and human disease. Nat Rev Genet 6, 597–610 (2005).

2. Bird, A. DNA methylation patterns and epigenetic memory. Genes Dev. 16, 6–21 (2002).

3. Peterson, S. N. & Reich, N. O. Competitive Lrp and Dam Assembly at the pap Regulatory Region: Implications for Mechanisms of Epigenetic Regulation. J. Mol. Biol. 383, 92–105 (2008).

4. Flusberg, B. a et al. Direct detection of DNA methylation during single-molecule, real-time sequencing. Nat. Methods 7, 461–5 (2010).

5. Frommer, M. et al. A genomic sequencing protocol that yields a positive display of 5-methylcytosine residues in individual DNA strands. Proc. Natl. Acad. Sci. U. S. A. 89, 1827–31 (1992).

6. Ip, C. L. C. et al. MinION Analysis and Reference Consortium: Phase 1 data release and analysis. F1000Research (2015). doi:10.12688/f1000research.7201.1

7. Schreiber, J. & Karplus, K. Analysis of Nanopore Data using Hidden Markov Models. Bioinformatics 1–7 (2015). doi:10.1093/bioinformatics/btv046

8. Timp, W., Comer, J. & Aksimentiev, A. DNA base-calling from a nanopore using a viterbi algorithm. Biophys. J. 102, L37–L39 (2012).

9. Jain, M. et al. Improved data analysis for the MinION nanopore sequencer. Nat. Methods (2015). doi:10.1038/nmeth.3290

10. Loman, N. J., Quick, J. & Simpson, J. T. A complete bacterial genome assembled de novo using only nanopore sequencing data. (2015).

11. Szalay, T. & Golovchenko, J. a. De novo sequencing and variant calling with nanopores using PoreSeq. Nat. Biotechnol. 33, 1–7 (2015).

12. Durbin, R., Eddy, S., Krogh, A. & Analysis, B. S. Biological Sequence Analysis: Probabilistic Models of Proteins and Nucleic Acids.

13. Wescoe, Z. L., Schreiber, J. & Akeson, M. Nanopores discriminate among five C5-cytosine variants in DNA. J. Am. Chem. Soc. 136, 16582–7 (2014).

14. Laszlo, A. H. et al. Detection and mapping of 5-methylcytosine and 5-hydroxymethylcytosine with nanopore MspA. Proc. Natl. Acad. Sci. U. S. A. (2013). doi:10.1073/pnas.1310240110

15. Teh, Y. W., Jordan, M. I., Beal, M. J. & Blei, D. M. Hierarchical Dirichlet Processes. J. Am. Stat. Assoc. 101, 1566–1581 (2006).

16. Schreiber, J. et al. Error rates for nanopore discrimination among cytosine, methylcytosine, and hydroxymethylcytosine along individual DNA strands. Proc. Natl. Acad. Sci. U. S. A. (2013). doi:10.1073/pnas.1310615110

17. Li, H. & Durbin, R. Fast and accurate short read alignment with Burrows-Wheeler transform. Bioinformatics 25, 1754–1760 (2009).

18. Paten, B., Herrero, J., Beal, K. & Birney, E. Sequence progressive alignment, a framework for practical large-scale probabilistic consistency alignment. Bioinformatics 25, 295–301 (2009).

19. Ferguson, T. S. A Bayesian Analysis of Some Nonparametric Problems. Ann. Stat. 1, 209–230 (1973).

## References

[1] Paten, B., Herrero, J., Beal, K., Fitzgerald, S. & Birney, E. Enredo and Pecan: genome-wide mammalian consistency-based multiple alignment with paralogs. Genome Research 18, 1814–1828 (2008).

[2] Teh, Y. W., Jordan, M. I., Beal, M. J. & Blei, D. M. Hierarchical Dirichlet Processes. Journal of the American Statistical Association 101, 1566–1581 (2006).

[3] Escobar, M. D. & West, M. Bayesian Density Estimation and Inference Using Mixtures. Journal of the American Statistical Association 90, 577–588 (1995).

